# Copy number variants (CNV) interfere with human Accelerated Regions (HARs) to cause neurodevelopmental disorders: a large-scale cohort study

**DOI:** 10.64898/2025.12.29.696895

**Authors:** Maria E. Iuditskaia, Ekaterina V. Semina, Elizaveta D. Pankratova, Oksana S. Kurinnaia, Elizaveta V. Guldan, Elvin H. Abasov, Ivan Y. Iourov, Yulia A. Chaika

## Abstract

Neurodevelopmental disorders (NDDs) represent a heterogeneous group of conditions affecting the central nervous system, the etiology of which involves a wide range of factors, including genetic contributors. A proportion of this genetic component is mediated by copy number variations (CNVs) - deletions and duplications of genomic segments that are associated with a broad spectrum of neurodevelopmental and neuropsychiatric phenotypes. In recent years, there has also been growing interest in studying Human Accelerated Regions (HARs) in relation to these disorders. HARs are highly conserved genomic elements across mammals that have accumulated human-specific sequence changes. These regions are thought to contribute both to the evolution of higher cognitive functions in humans and, paradoxically, to an increased susceptibility to NDDs. Thus, both HARs and CNVs are expected to play key roles in the genetic architecture and pathogenesis of NDDs. However, despite active research on HARs and CNVs separately, integrative studies examining copy-number–driven disruptions specifically within HARs in patients with neurodevelopmental disorders have been extremely limited.

This is the first study to assess the contribution of CNVs affecting HARs in 821 children with NDDs. Our bioinformatic evaluation of CNV profiles in this clinical cohort demonstrated that HARs are indeed susceptible to structural alterations in the form of deletions and duplications, supporting the notion that CNVs have shaped the evolution of these regions. Our findings further indicate that HARs, overall, are less frequently affected by duplications compared to other genomic regions. We identified three recurrent HAR loci most strongly associated with neurodevelopmental impairment: HAR_Merge50-02702, HAR_Merge50-02080, and HAR_Merge50-02689. Additionally, we found that HARs disrupted by CNVs form coordinated functional clusters corresponding to genomic regions that harbor extremely rare or entirely absent CNVs in the general population (e.g., HARs associated with *RBFOX1, DOC2B, ZMYND11, AFF2, OPHN1*). These same genes exhibit a high haploinsufficiency index (HI = 3) according to ClinGen, highlighting their dosage sensitivity. Analysis of the most frequently affected HARs confirmed that their associated genes are significantly enriched for core NDD phenotypes. The absence of a direct correlation with genomic instability suggests that HARs represent functionally vulnerable elements and that compensatory mechanisms may restrict the accumulation of additional structural damage in these regions. Taken together, our study reinforces the importance of HARs in the evolution and functioning of the human central nervous system and provides new evidence supporting the functional relevance of HAR-associated disruptions in neurodevelopmental disorders. We anticipate that future research will further elucidate the complex regulatory processes within the genome and help identify specific HAR regions that may be informative for diagnostic purposes and the development of targeted therapeutic approaches.

## 1. Introduction

Across the ∼8 million years of human genomic evolution since the divergence from our closest living ancestor, the human genome has undergone numerous changes that contributed to the emergence of uniquely human cognitive and behavioral traits. Developments of higher cognitive abilities—such as memory, language, and complex reasoning—are thought to be linked to the expansion and increasing structural complexity of the cerebral cortex, which enabled the formation of more elaborate neural architectures [1–3]. Because the human and chimpanzee genomes demonstrate extraordinarily similarity and protein-coding genes show relatively few human-specific changes [4], it has been proposed that the genetic basis of these differences may lie primarily in noncoding regulatory regions, such as Human Accelerated Regions (HARs) [5–8]. Accordingly, bioinformatic approaches in 2006 led to the discovery of HARs [9], which may represent a major driver of human evolution. HARs are evolutionarily conserved genomic elements across mammals that have accumulated exceptionally rapid DNA sequence changes since the human–chimpanzee split. Subsequent studies expanded the original list [6,8,10–12] and today approximately 3,000 HARs have been identified [13,14].

The majority of HARs (∼96%) are noncoding and located within intronic or intergenic regions [15], with nearly half functioning as tissue-specific transcriptional enhancers involved in regulating genes essential for nervous system development [7,15–19]. Moreover, many HARs act as enhancers during embryonic neurodevelopment [20,21], and in brain structures that exhibit human-specific morphological features [22,23]. Importantly, enhancer activity of certain HARs differs markedly from that of chimpanzee orthologs [16,24–26] with some HARs showing substantially increased activity in humans [15], suggesting that they may have contributed to the emergence of human-specific brain development.

Of particular interest is the strong enrichment of HARs near genes that play critical roles in neurodevelopmental processes, including neural progenitor proliferation, neuronal migration, and synapse formation [13,27]. Disruption of these regulatory regions has been hypothesized to alter the expression of key genes implicated in autism spectrum disorders (ASD), schizophrenia, and other psychiatric conditions, underscoring their potential clinical relevance [14,28]. Consequently, interest in the contribution of HARs to the pathogenesis of NDDs has grown considerably in recent years [29,30]. Most existing studies have focused on HAR sequence mutations (primarily single-nucleotide changes) in schizophrenia [31–36], whereas NDDs as a broader category remain understudied, with relatively few publications available [3,30,37]. Additional research has also examined HAR involvement in other psychiatric and neurodegenerative disorders [36–39].

NDDs are a heterogeneous group of conditions involving abnormalities in the formation, maturation, and function of the nervous system, manifesting as developmental delay, cognitive deficits, epilepsy, and behavioral impairments. Their etiology is typically complex, but genetic factors, including rare structural variants, are considered major contributors [40–42]. Copy number variations (CNVs)—deletions and duplications of genomic segments—play a particularly important role in the genetic architecture of NDDs and are associated with a broad spectrum of neurodevelopmental and neuropsychiatric phenotypes [40,41,43]. CNVs may affect both coding genes and distant regulatory elements, including enhancers and highly conserved noncoding sequences active during brain development. Accumulating evidence suggests that disruptions in such regulatory regions can contribute to severe NDD phenotypes [43–45].

Despite active research on HARs and CNVs, comprehensive analysis of CNV-driven disruptions within HARs in patients with neurodevelopmental disorders has been almost missed. Only a single study to date has reported an association between HARs and CNVs, demonstrating that de novo CNVs affecting HARs or HAR-containing genes may account for up to 1.9% of ASD cases [14]. Given that HARs frequently function as enhancers in neuronal sells, it is reasonable to hypothesize that structural variants (CNVs) involving these evolutionarily significant regulatory regions could have particularly severe consequences for brain development and may contribute to psychiatric and neurodevelopmental pathology. Alterations in the structure or dosage of a HAR enhancer may dysregulate networks of downstream target genes involved in neurogenesis and synaptic function, representing a plausible pathological mechanism.

In this study, we performed a bioinformatic analysis of molecular karyotyping data from 821 individuals with NDDs to identify and characterize HAR regions affected by CNVs. As no prior study has systematically investigated the interplay between HARs and CNVs in this context, our work provides a novel contribution to the field. Here, we assessed the frequency of HAR-disrupting CNVs, explored their evolutionary significance, conducted cluster analysis of co-affected HAR regions, and examined the contribution of key disrupted HARs to relevant phenotypes. This represents the first comprehensive study aimed at understanding the relationship between HARs and CNVs in NDDs. Although this is a pilot study, we anticipate that future research will shed further light on the complex mechanisms underlying these disorders.

## 2. Results and discussion

### Analysis of HAR Occurrence in Neurodevelopmental Disorders

We analyzed molecular karyotyping data (the full set of CNVs) from 821 patients with neurodevelopmental disorders. We found that a substantial proportion of the cohort—755 patients—carried CNVs overlapping HAR regions, indicating that HARs are susceptible to structural variation, which may have not only pathogenic but also evolutionary relevance. In total, 6,090 CNVs affecting HARs were identified, including 3,188 deletions (52.35%) and 2,902 duplications (47.65%) (Supplementary Table 1).

Within the cohort, we identified the three most frequently affected HARs: HAR_Merge50-02702 (7.9%), HAR_Merge50-02080 (∼4.0%), and HAR_Merge50-02689 (∼2.9%). These HARs have not been previously highlighted in the literature and have not been subjected to detailed analysis. Less frequent—but still notable—HARs included HAR_Merge50-01379 (1.9%), HAR_Merge50-01157 (1.2%), and HAR_Merge50-01486 (1.1%).

**HAR_Merge50-02702** is associated with the genes *PLS3* and *AGTR2* [14]. *PLS3* is a Ca²⁺-dependent F-actin–bundling protein that influences the G/F-actin ratio and plays an important role in the formation and maintenance of neuromuscular synapses [46]. Actin dynamics are essential for axon development, cell polarity, migration, axonal outgrowth, receptor trafficking (e.g., *TrkB*), and neuronal plasticity. Its expression markedly increases during the development and maturation of motor neurons, highlighting the importance of *PLS3* in these processes [47]. *AGTR2* encodes the angiotensin II type-1 receptor (*AT1R*), which promotes oxidative stress, apoptosis, and neuroinflammation leading to neurodegeneration, and may play an important role in maintaining human brain function [48].

**HAR_Merge50-02080** is associated with *BTNL3* and *BTNL8*, which are TCR-dependent regulators of human intestinal γδ T cells [49].

**HAR_Merge50-02689** affects the gene *EDA*, mutations in which impair the development of multiple ectodermal structures, including hair, sweat glands, and teeth [50]. The receptor for *EDA*, *EDAR*, is expressed in the neural crest, giving rise to cranial ganglia and glia. TNFR-pathway signaling typically activates the transcription factor *NF-κB* [51], which is involved in neuronal survival, synaptic plasticity, and neurological disease [52,53]. Thus, *EDA*-mediated activation of *NF-κB* may modulate gene expression programs related to synaptic transmission and neuroinflammation.

**HAR_Merge50-01379** is associated with *ERBB4*, a receptor tyrosine kinase of the ErbB family. It is a key component of the Neuregulin–ErbB4 pathway, regulating the development and function of GABAergic interneurons, synaptic plasticity, and excitatory/inhibitory balance. *ERBB4* participates in cell proliferation, differentiation, migration, angiogenesis, and apoptosis and plays a central role in interneuron maturation. It is strongly associated with schizophrenia and cognitive impairment[54–56].

**HAR_Merge50-01157** is associated with the genes *ASXL2*, *KIF3C*, and *GAREM2* (previous HGNC Symbols: *FAM59B*)*. ASXL2* is an essential epigenetic regulator required for proper chromatin state and gene expression [57]. Germline de novo truncating *ASXL2* variants cause Shashi–Pena syndrome, characterized by developmental delay, glabellar nevus flammeus, hypotonia, and cardiac abnormalities [58,59]. *KIF3C* encodes a motor protein involved in axonal transport and neurite outgrowth and plays a key role in neuronal maturation [60,61]. *GAREM2* encodes an adaptor protein in the MAPK pathway, is predominantly expressed in the brain, contributes to cognition and behavior, and may be involved in neurodegeneration. *GAREM2*-knockout mice display altered anxiety and exploratory behavior [62].

**HAR_Merge50-01486** contains the genes *CYYR1* and *ADAMTS1*. *CYYR1* has been proposed as a potential biomarker for borderline personality disorder [63]. *ADAMTS1* belongs to the ADAMTS family of extracellular metalloproteinases; in the CNS, ADAMTS proteins remodel the extracellular matrix, play essential roles in neuroplasticity, and contribute to neurological pathologies such as Alzheimer’s disease [64].

We identified HARs that regulate coordinated molecular programs essential for CNS development and function. Disruptions in HAR_Merge50-02702 may influence axonal growth, synaptic plasticity, and receptor trafficking (e.g., *TrkB*), suggesting that CNVs in this HAR could underlie disturbances in neuronal network formation and stability. The second most frequent HAR, HAR_Merge50-02080, is associated with *BTNL* family genes involved in γδ T-cell regulation; although their brain-specific functions are less understood, this suggests a potential link between immune regulation and neurodevelopment. The third region, HAR_Merge50-02689, affects the *EDA/EDAR* pathway, which activates *NF-κB*—a key regulator of neuronal survival, synaptic plasticity, and neuroinflammation. Analysis of less frequent but functionally meaningful HARs (HAR_Merge50-01157, HAR_Merge50-01379, HAR_Merge50-01486) revealed disruptions in genes involved in essential cellular processes: epigenetic control (*ASXL2*), intracellular transport (*KIF3C*), MAPK-signaling (*GAREM2*), and extracellular matrix remodeling (*ADAMTS1*).

Our findings support the hypothesis that CNVs affecting HAR regions in individuals with NDDs are not merely byproducts of genomic instability, but events that directly perturb finely tuned evolutionary innovations of the human genome. These disruptions involve coordinated functional modules regulating core neurodevelopmental processes—from cytoskeletal organization and synaptogenesis to epigenetic programming and intercellular communication. A notable pattern is the preferential disruption of HARs associated with genes integrating external signals (growth factors, neurotransmitters, immune stimuli) into long-term neuronal changes. Thus, our results establish a direct link between variation in evolutionarily significant noncoding regions and the clinical phenotype of NDDs. This supports the idea that HARs contributed to the evolution of the human brain, and that their disruption in NDDs is functionally meaningful, affecting regulatory programs of multiple genes—likely through their enhancer activity. Further molecular studies will be needed to validate these effects.

To assess the extent to which HAR regions are affected by CNVs, we performed an independent permutation analysis (1000 permutations) evaluating whether the observed number of CNV–HAR intersections exceeds that expected under a random genomic distribution. We found that deletions (loss CNVs) showed a weak but non-significant deviation from expectation (obs = 2218; mean = 2266; z = –1.10; p > 0.05), whereas duplications (gain CNVs) demonstrated a pronounced depletion (obs = 1764; mean = 2174.7; z = –10.13; p < 0.001). When considering all CNVs together, we observed a strong and statistically significant depletion of CNV–HAR intersections (obs = 3982; mean = 4644.1; z = –7.54; p < 0.001) (Fig. 1E).

**Figure 1.**
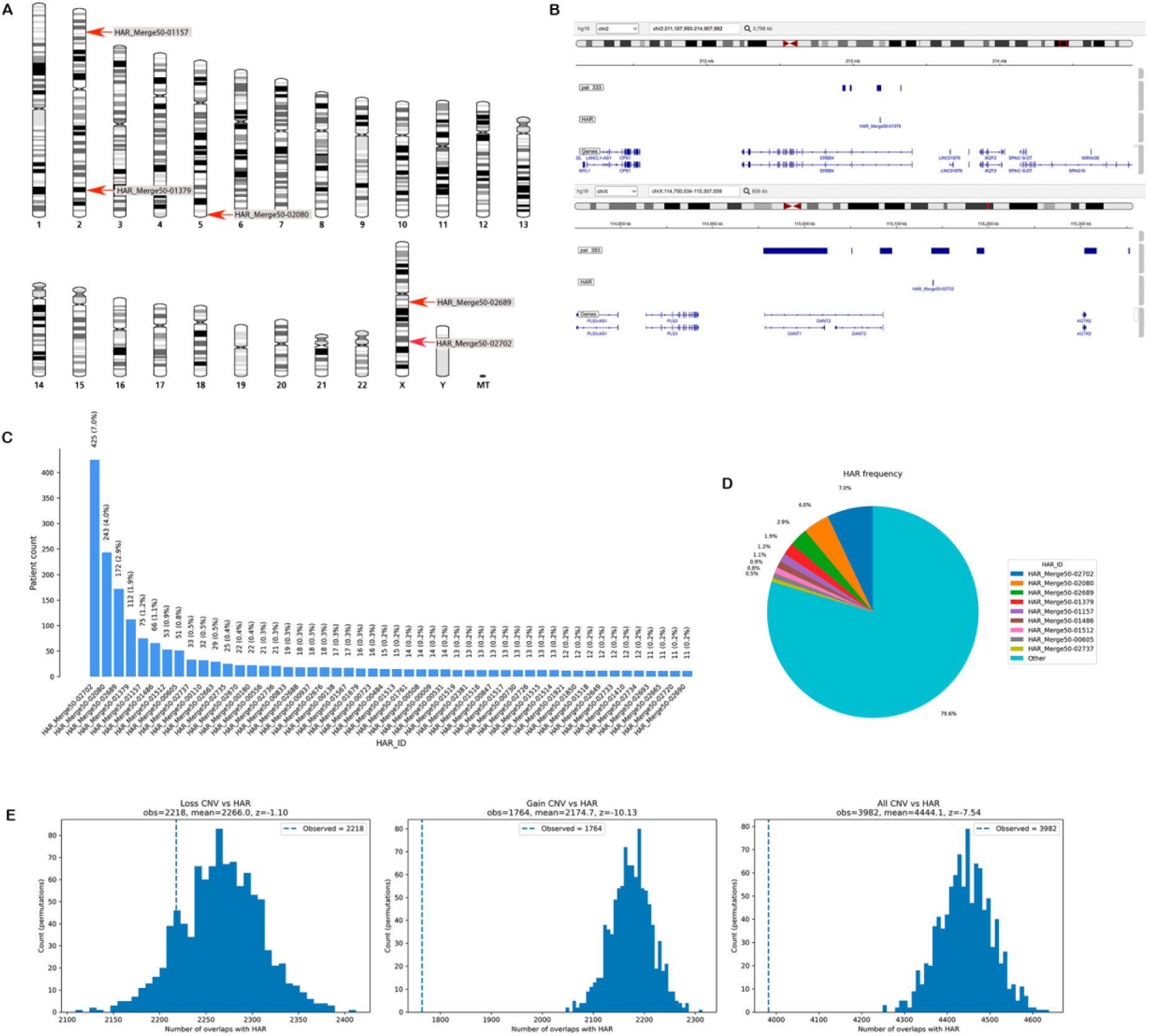
Genomic landscape of CNV–HAR intersections in patients with neurodevelopmental disorders. **A** – Genomic locations of the most frequently affected HAR regions in the cohort. Red arrows indicate HAR_Merge50-01157 (chr2), HAR_Merge50-01379 (chr2), HAR_Merge50-02080 (chr5), HAR_Merge50-02689 (chrX), and HAR_Merge50-02702 (chrX). **B** – Visual annotation of selected recurrent HAR loci disrupted by CNVs. **C** – Frequency distribution of the 50 HARs affected by CNVs in the analyzed cohort. Histogram showing the distribution of CNV–HAR intersection counts; lower-frequency HARs were omitted for clarity. Each bar corresponds to an individual HAR_ID, and the Y-axis indicates the number of patients carrying a CNV affecting the given HAR. The most frequently observed regions are HAR_Merge50-02702, HAR_Merge50-02080, and HAR_Merge50-02689. **D** – Relative contribution of the most frequent HARs to the total number of CNV–HAR intersections. Pie chart illustrating the contribution of HAR regions (n = 9, total intersections = 6035) to the overall number of CNV overlaps. HARs were selected as loci with the highest number of observations among all 2737 regions from the Doan et al. (2016) dataset, representing the top HARs by frequency. HAR_Merge50-02702 accounts for ∼7.9% of all intersections, followed by HAR_Merge50-02080 (∼4.0%), HAR_Merge50-02689 (∼2.9%), and others. The “Other” category includes all remaining low-frequency HARs. **E** – Permutation test assessing the enrichment of CNVs within HAR regions. Each panel shows the distribution of overlap counts obtained from randomized CNV sets (histogram) and the observed value (vertical dashed line).

The strong depletion of duplications overlapping HARs is consistent with the view that HARs are functionally important and are subject to negative selection against structural variants that increase copy number of these regulatory element [65]. While deletions exhibit only a weak and statistically non-significant trend toward depletion, duplications show an approximately 19% deficit relative to the null model. This may reflect stronger evolutionary constraints on duplications in regulatory elements involved in development and brain function compared with deletions of similar length and genomic distribution [66,67]. Although deletions of conserved regulatory elements are often pathogenic (haploinsufficient) and eliminated by selection [66], in our analysis we observe the opposite pattern, which may indicate either a minimal functional impact of HAR deletions or, conversely, the emergence of early adaptive processes in these regions, or a potential contribution to disease phenotypes. However, when deletions and duplications are analyzed jointly, the overall depletion strongly supports the presence of negative selection acting specifically against CNV disruption of HAR regions [68], reinforcing their functional significance and essential role in human CNS development [14].

### Patterns of HAR Disruption and Cluster Analysis

To evaluate patterns of co-occurring disruptions in Human Accelerated Regions (HARs) within the cohort, we selected the 50 most frequently affected HARs (see Supplementary Table 2). For this subset, two pairwise relationship matrices were constructed (Fig. 2):

**A** – a co-occurrence matrix representing the absolute number of patients in whom two HARs were disrupted simultaneously;

**B** – a Jaccard similarity matrix providing a normalized measure of overlap, allowing the detection of proportionally related HARs even when their absolute frequencies are low.

The co-occurrence matrix (Fig. 2A) identified groups of HARs that tend to be affected in the same individuals, which may reflect shared genomic rearrangements or coordinated regulatory interactions. In contrast, the Jaccard similarity matrix (Fig. 2B) highlighted pairs of HARs with a high degree of proportional overlap, even when co-occurrence counts were low, indicating potential structural or functional relationships between the corresponding genomic regions. Hierarchical clustering of the Jaccard matrix (Fig. 2B) revealed four robust HAR clusters (see Supplementary Table 3). Analysis of intracluster similarity showed that in clusters 1–3, Jaccard coefficients exceeded 0.1 for most HAR pairs (range 0.1 to 1), indicating strong coherence and functional relatedness of elements within these modules (Fig. 2B). In contrast, cluster 4 exhibited predominantly low Jaccard values (most pairs < 0.1), suggesting a more heterogeneous structure and was therefore excluded from further analysis.

**Figure 2.**
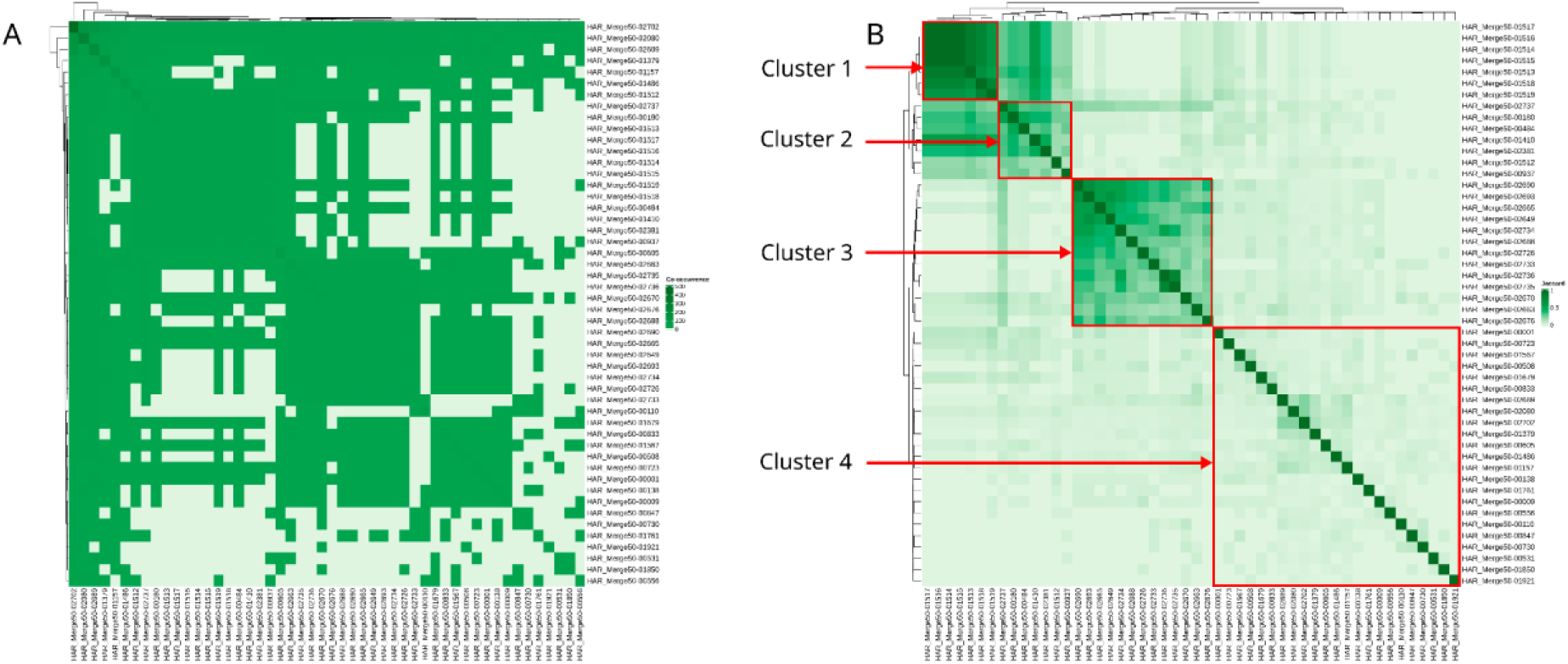
Heatmaps of co-occurrence and Jaccard similarity among the 50 most frequent HARs. **A** – Co-occurrence matrix showing the absolute number of patients carrying each HAR pair. **B** – Jaccard similarity matrix representing the normalized overlap between HARs. Darker shades of green indicate higher values of joint appearance or similarity.

### Cluster 1

Cluster 1 includes HARs associated with the genes *CHL1* and *CNTN6*, both located in the 3p26.3 locus. Both genes belong to the family of neural cell adhesion molecules and play essential roles in axonal growth and guidance, synapse formation, and cortical development [14,69,70]. The high degree of intracluster similarity suggests that the HARs within this module may reflect a shared regulatory mechanism influencing the function of neural cell adhesion systems. Moreover, because multiple HARs overlap *CHL1* and *CNTN6*, this may indicate particularly fine and strong regulatory control over these genes; consequently, disruption of several HARs simultaneously is likely to exert stronger effects at the molecular level.

### Cluster 2

Cluster 2 includes HARs and associated with the genes *CHL1*, *DOC2B*, *ERICH1-AS1*, *PSMF1*, *RSPO4*, *SPRY3*, *ZMYND11*, and *ZNF891*. This cluster represents a functional module whose elements are located on different chromosomes but are united by shared pathways and regulatory processes. The protein encoded by *RSPO4* is an activator of Wnt signaling [71], whereas *SPRY3* is an inhibitor of the FGF/RTK pathway [72]. Knockout or overexpression of *SPRY3* affects neurite growth and branching [73]. *DOC2B* participates in calcium-dependent regulation of synaptic neurotransmitter release [74], and *ZMYND11* is an epigenetic modifier that influences transcriptional programs of neuronal differentiation and is associated with intellectual developmental disorder [75–77]. Thus, HARs in Cluster 2 may participate in regulating neurodevelopment, particularly through modulation of Wnt/FGF signaling and synaptic processes essential for CNS function. The genes *RSPO4*, *SPRY3*, and *CHL1* act as regulators of fundamental pathways (Wnt, FGF, RTK), which control neurogenesis, neuronal migration, axon extension, and synaptogenesis. Evolutionary changes in HARs may have fine-tuned the balance and precision of these signals, contributing to the increasing complexity of neural circuitry. Disruption of this balance—such as that observed with *SPRY3* in autism [73] —underlies many neurodevelopmental disorders. The genes in this cluster operate at molecular and cellular levels: from immediate neurotransmitter release (*DOC2B* [74]) to long-term epigenetic regulation of hundreds of genes determining neuronal fate (*ZMYND11*). *ZMYND11* is particularly notable because the mutations cause an autosomal dominant syndrome, associated with intellectual disability and behavioral abnormalities [75].

Taken together, HARs in Cluster 2 likely represent elements of an advanced regulatory network that, enabled higher coordination between, during human evolution: growth signaling pathways (Wnt, FGF), cell adhesion and communication (*CHL1*), synaptic precision (*DOC2B*), epigenetic programs of neuronal specialization (*ZMYND11*). This enhanced coordination may have contributed to the complexity of cortical structures and cognitive functions. Accordingly, disruption of these finely tuned evolutionary sequences (HARs) or their associated genes increases the risk of neurodevelopmental disorders such as autism spectrum disorders, intellectual disability, and syndromic developmental delay—consistent with the characteristics of our cohort.

### Cluster 3

Cluster 3 includes HARs associated with the genes *AFF2*, *DACH2*, *DMD*, *FMR1NB*, *HSFX1*, *IL1RAPL1*, *MAGEA11*, *MXRA5*, *OPHN1*, *PRKX*, *SPANXC*, *TTC3P1*, *ZDHHC15*, *LINC00894*, and *MIR2114*, most of which are located on the chromosome X.

The gene *DMD* encodes dystrophin, mutations in which cause Duchenne muscular dystrophy [78]. Additionally, *DMD* is associated with a spectrum of brain abnormalities ranging from mild cognitive impairment (MCI) to neuronal migration disorders [79–81]. *IL1RAPL1* and the nearby HAR regions are associated with intellectual disability and autism [37]*. RGS3* and *IL1RAPL1* missense variants implicate defective neurotransmission in early-onset inherited schizophrenia [82]. *AFF2* is involved in the FRAXE syndrome, which is associated with intellectual disability, obsessive–compulsive disorder, and primary ovarian insufficiency [83]. *OPHN1* is linked to X-linked intellectual disability [84,85].

Cluster 3 represents a functionally integrated X-linked module that likely underwent coordinated evolutionary shaping under the influence of HARs. Its primary biological role is the regulation of synaptic architecture and plasticity. The localization of most genes on the X chromosome has two key implications. 1. Evolutionary “laboratory”: The X chromosome, particularly in placental mammals, is enriched for genes associated with brain development and cognitive functions. This makes it a “hotspot” for the evolution of traits related to intelligence and social behavior. HARs in this region may have fine-tuned the dosage and expression patterns of an entire set of neuronal genes simultaneously [37,85]. 2. Clinical vulnerability: Mutations in X-linked genes manifest phenotypically in hemizygous males, contributing to the male bias in autism spectrum disorders, ADHD (attention deficit hyperactivity disorder), and intellectual disability. Cluster 3 includes several established causal genes for these conditions (*IL1RAPL1*, *OPHN1*, *ZDHHC15*, *AFF2*). The genes in this cluster are not specifically brain-related; these are associated with a specific neuronal structure—the synapse. *DMD* stabilizes the membrane and clusters receptors [79,80]; *OPHN1* (via Rho GTPase) and *PRKX* (via PKA) regulate the cytoskeleton and synaptic strength [85,86]; *ZDHHC15* palmitoylates key synaptic proteins such as PSD-95, regulating their stability [87]; *IL1RAPL1* and *AFF2* couple synaptic activity to long-term gene expression changes [37,88,89]. HARs in Cluster 3 likely evolved to provide coordinated regulation of this synaptic ensemble on the X chromosome. Their modification may have optimized postsynaptic transmission—one of the molecular foundations for the increased complexity of neural circuits underlying learning, memory, and social cognition. Disruption of this fine-tuned system (via mutations in the genes themselves or in their HAR regulators) predictably leads to a spectrum of neurodevelopmental disorders with strong male predominance.

Our analysis of genomic rearrangements within Human Accelerated Regions (HARs) in a cohort of children with neurodevelopmental disorders identified three functionally meaningful clusters representing coordinated regulatory modules associated with the pathogenesis of neurodevelopmental dysfunction. Cluster 1, affecting the 3p26.3 locus and the neuronal adhesion genes *CHL1* and *CNTN6*, highlights disruptions in core mechanisms of axonal growth and neuronal wiring. Cluster 2, a cross-chromosomal module involving *CHL1*, *DOC2B*, *RSPO4*, *SPRY3*, *ZMYND11*, and others, reflects a broader systemic imbalance affecting key developmental signaling pathways (Wnt/FGF), synaptic transmission, and epigenetic programs of neuronal specialization—features characteristic of severe and syndromic neurodevelopmental disorders. Cluster 3, enriched for X-linked genes (*DMD*, *IL1RAPL1*, *AFF2*, *OPHN1*, *ZDHHC15*), forms a clear molecular substrate linking disruptions in synaptic architecture and plasticity to clinical manifestations such as intellectual disability, autism spectrum disorders, and epilepsy. Together, these findings indicate that HAR disruptions are not random but preferentially affect evolutionarily refined regulatory ensembles that coordinate complex neurodevelopmental processes. Perturbation of these finely balanced modules directly contributes to CNS pathology, underscoring the functional importance of HARs and opening new avenues for understanding their roles in the diagnosis and pathogenesis of neurodevelopmental disorders.

### Analysis of the Uniqueness of CNV–HAR Frequencies in Our Cohort Compared with Population Databases

To evaluate the uniqueness of alterations in the top 50 HARs within our cohort, we compared the observed frequencies of HAR-disrupting CNVs in our patients with data from publicly available population databases (ClinGen [90], DECIPHER [91] и DGV [92]). Numerical values are summarized in Supplementary Table 5. Each point on the scatter plot (Fig. 3A) represents a HAR locus, where the x-axis shows the deviation in duplication frequency (ΔGain) and the y-axis shows the deviation in deletion frequency (ΔLoss) relative to database-derived background rates.

**Figure 3.**
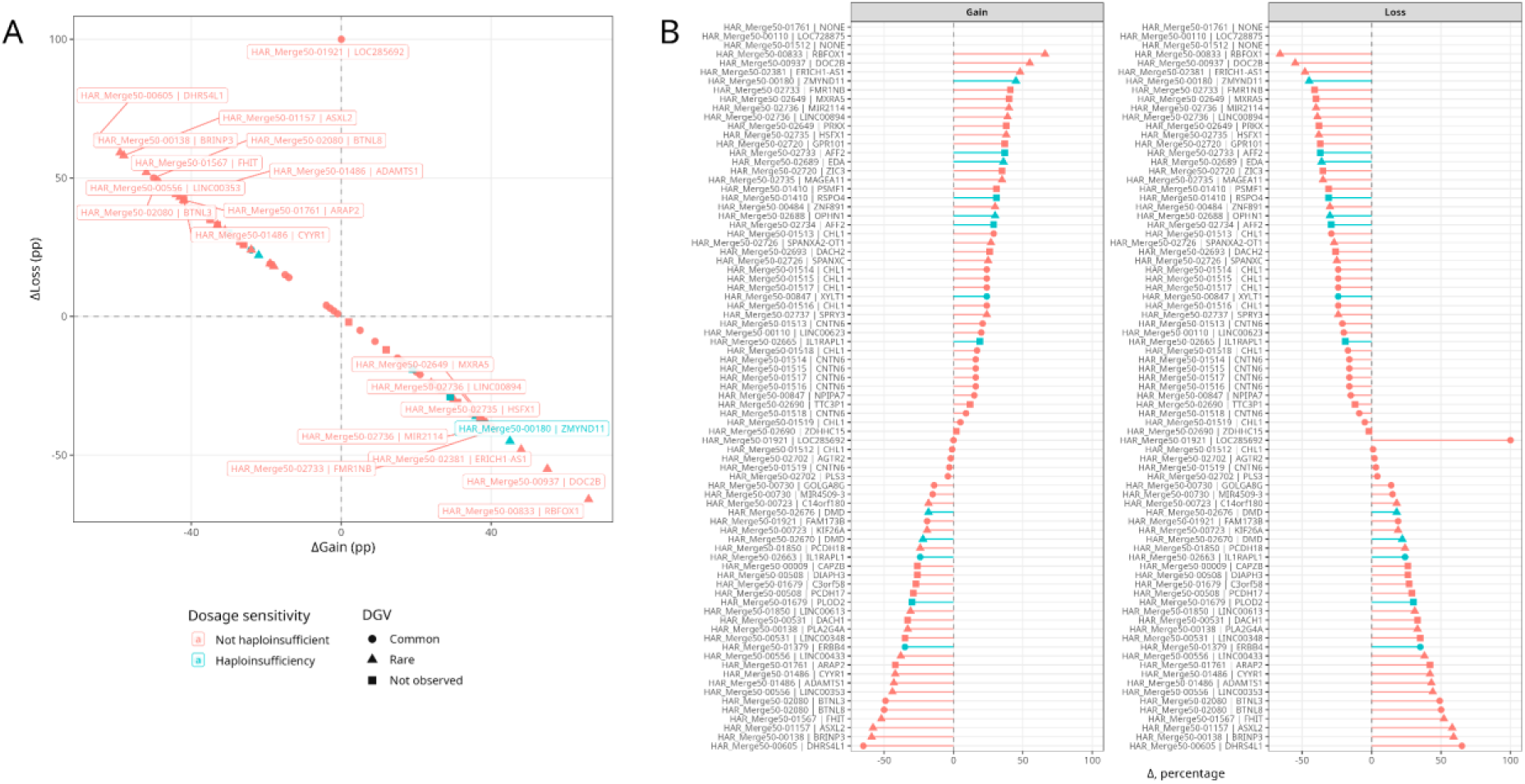
Comparative analysis of CNV patterns across Human Accelerated Regions (HARs). **A** - Scatter plot showing differences between observed CNV frequencies in the studied cohort and reference frequencies reported in DGV for copy-number gains (ΔGain, x-axis) and losses (ΔLoss, y-axis). Each point corresponds to a HAR locus. Point color reflects ClinGen dosage sensitivity status (red — no evidence of dosage sensitivity, ClinGen = 0; cyan — dosage-sensitive, ClinGen = 1). Symbol shape indicates DGV population frequency class (circle — common, triangle — rare, square — not observed). **B** - Ranked distributions of ΔGain (left) and ΔLoss (right) values for all HARs. Points and symbols follow the same ClinGen and DGV conventions as in panel A. Positive values indicate enrichment in the studied cohort relative to population references, while negative values represent lower-than-expected frequencies.

For most HARs, the deviation from reference values is substantial (Fig. 3A), suggesting their potential role in the pathogenesis of NDDs. In particular, HAR_Merge50-00833 (*RBFOX1*), HAR_Merge50-00937 (*DOC2B*), HAR_Merge50-02381 (*ERICH1-AS1*), HAR_Merge50-00180 (*ZMYND11*), HAR_Merge50-02733 (*FMR1NB*), HAR_Merge50-02649 (*MXRA5*), HAR_Merge50-02736 (*MIR2114*, *LINC00894*), HAR_Merge50-02735 (*HSFX1*), HAR_Merge50-02720 (*GPR101*), and HAR_Merge50-02733 (*AFF2*) exhibited markedly elevated duplication frequencies (ΔGain > 0) compared with population baselines. Conversely, HAR_Merge50-00605 (*DHRS4L1*), HAR_Merge50-00138 (*BRINP3*), HAR_Merge50-01567 (*FHIT*), HAR_Merge50-00556 (*LINC00353*), HAR_Merge50-02080 (*BTNL3*, *BTNL8*), and HAR_Merge50-01157 (*ASXL2*) demonstrated increased deletion frequencies. Some HARs display asymmetric profiles (Gain↑ with Loss↓, or vice versa). A cluster of loci shows strongly positive ΔGain values with ΔLoss near or below zero—an imbalance favoring duplications—typical for HARs associated with *RBFOX1*, *DOC2B*, and *ZMYND11*. In contrast, *BTNL3/BTNL8*, *FHIT*, *BRINP3*, and *ASXL2* exhibit predominantly deletions.

When comparing our CNV frequencies with the DGV dataset (categorizing variants as rare/not observed), many HARs with the strongest ΔGain/ΔLoss deviations correspond to loci where population CNVs are either rare or entirely unreported. Among rare CNVs: HAR_Merge50-00833 (*RBFOX1*) shows rare CNVs associated with epilepsy, ASD, and other neurodevelopmental phenotypes; HAR_Merge50-00937 (*DOC2B*) involves a synaptic regulatory gene; HAR_Merge50-00180 (*ZMYND11*) shows rare CNVs despite the gene’s high haploinsufficiency index (HI = 3) in ClinGen; HAR_Merge50-02733 (*AFF2*), HAR_Merge50-02688 (*OPHN1*), HAR_Merge50-02676 (*DMD*), HAR_Merge50-02663 (*IL1RAPL1*) represent rare X-linked CNVs associated with intellectual disability and neurodevelopmental delay.

Not observed CNVs in DGV: HAR_Merge50-02733 (*FMR1NB*), HAR_Merge50-02649 (*MXRA5*), HAR_Merge50-02649 (*PRKX*), HAR_Merge50-02720 (*GPR101*), HAR_Merge50- 02733 (*AFF2*), HAR_Merge50-02720 (*ZIC3*), HAR_Merge50-01761 (*ARAP2*), HAR_Merge50-00531 (*LINC00348*), HAR_Merge50-00531 (*DACH1*), HAR_Merge50- 01679 (*PLOD2*). Thus, for many HARs interacting with key neurodevelopmental genes, the corresponding CNVs are either extremely rare or completely absent in DGV, yet show substantial frequency shifts in our cohort of children with CNS disorders.

Several HARs associated with dosage-sensitive genes show strong deviations in ΔGain/ΔLoss, are enriched for rare or unreported CNVs, and are linked to neurodevelopmental phenotypes in ClinGen/DECIPHER: *ZMYND11* (HI = 3): complex neurodevelopmental disorder, speech delay, intellectual disability, seizures; *DMD* (HI = 3): classic dosage-sensitive muscular dystrophy; in our data, the HAR near *DMD* deviates in ΔLoss; *OPHN1* (HI = 3): X-linked intellectual disability with cerebellar hypoplasia; *IL1RAPL1* (HI = 3): X-linked intellectual disability and ASD; *AFF2* and other X-linked dosage-sensitive genes.

These are precisely the loci where any additional CNV affecting a HAR enhancer is likely to influence gene expression and phenotype. In our analysis, structural alterations in these HARs occur at frequencies different from CNVs in the genes themselves (ΔGain/ΔLoss ≠ 0). This pattern is most pronounced for genes involved in splicing and synaptic function (*RBFOX1*, *DOC2B*), epigenetic regulation (*ZMYND11*, *ASXL2*), X-linked dosage-sensitive loci (*OPHN1*, *IL1RAPL1*, *AFF2*, *DMD*), and cell-adhesion molecules (*CNTN6*, *CHL1*), which are already implicated in 3p deletion syndrome and neurodevelopmental disorders.

The rarity of these CNVs in the general population (DGV) and their dosage sensitivity (ClinGen) reinforce the interpretation that the observed events are not “noise,” but may contribute directly to the phenotype of our CNS-affected cohort.

Furthermore, pathogenic CNVs cluster within functionally organized HAR modules. The largest deviations from population background (ΔGain/ΔLoss) and the rare/not-observed status are characteristic of HARs associated with genes belonging to specific regulatory clusters. This indicates that structural variants disrupt not random genomic points but coordinated regulatory ensembles (Clusters 1–3) controlling key neurodevelopmental processes: adhesion and migration (Cluster 1), intercellular signaling and synaptic function (Cluster 2), and X-linked synaptic architecture (Cluster 3).

The likely mechanism of pathogenicity is dosage sensitivity disruption. Many cluster genes (*ZMYND11*, *OPHN1*, *DMD*, *IL1RAPL1*) have high haploinsufficiency indices (HI = 3). Even subtle expression changes resulting from CNVs within associated HAR enhancers may produce clinically significant effects. Thus, the observed CNVs likely act as regulatory dosage alterations (enhanceropathies), consistent with severe phenotypes in cases with large structural rearrangements.

Finally, the rarity of HARs in the population strongly supports purifying selection and clinical relevance. The fact that CNVs linked to cluster-associated genes (*DOC2B*, *ZMYND11*, *AFF2*, etc.) are rare or absent in DGV indicates their deleterious nature. Their enrichment in our cohort of children with CNS disorders provides strong evidence for a causal contribution to pathology.

### Phenotypes of HARs and Their Associated Genes

We also analyzed genes associated with the top 50 human accelerated regions (HARs) using data from the DECIPHER database [91]. The aim was to determine which clinical diagnoses are linked to genes located near HARs and to identify shared phenotypic patterns (full list in Supplementary Table 5).

Each HAR was matched to its nearest or overlapping genes, after which clinical diagnoses recorded in DECIPHER (Morbid field) were extracted for these genes. Based on these annotations, we constructed a binary “gene–disease” matrix, where the presence of an association is shown in turquoise and its absence in gray (Fig. 4A).

**Figure 4.**
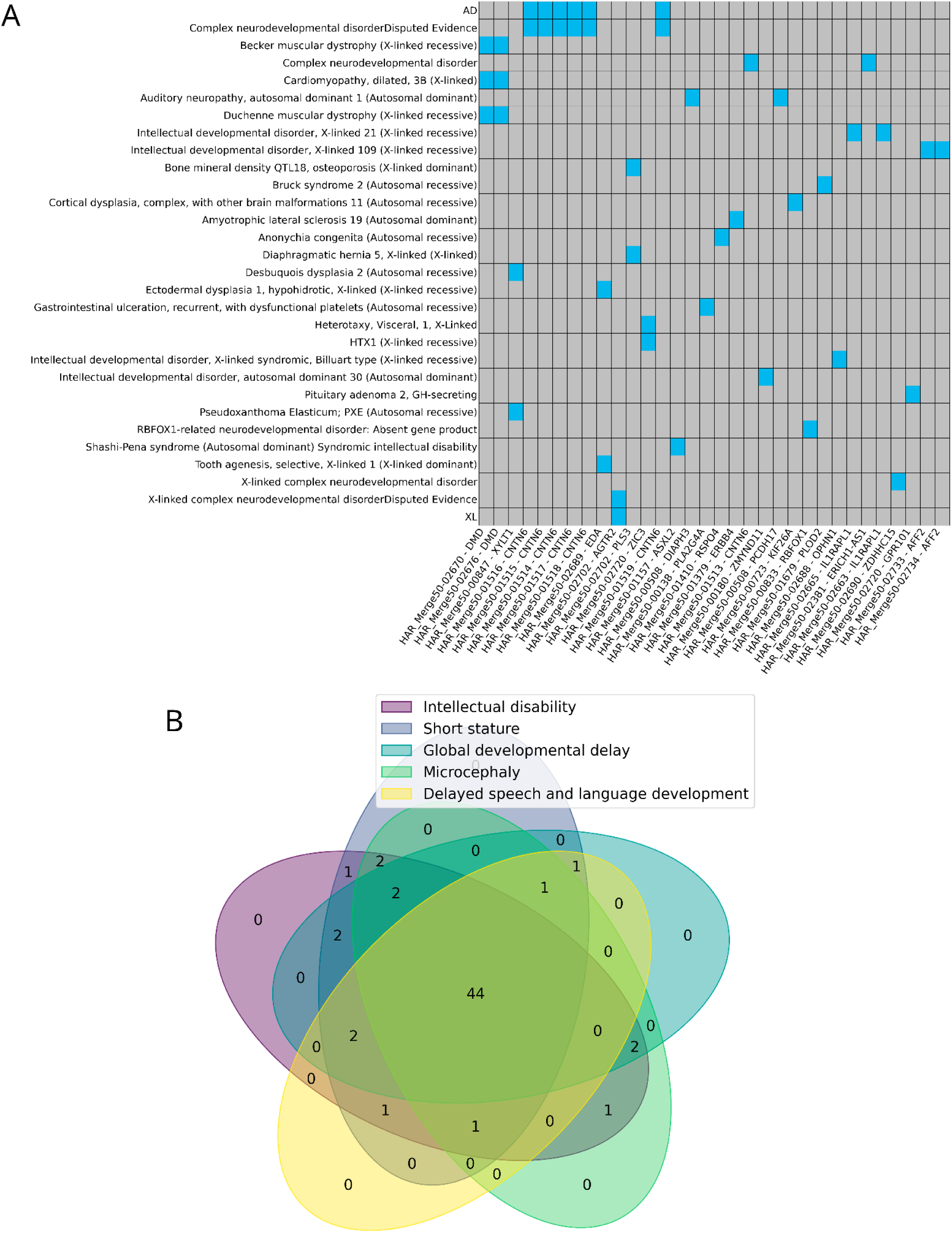
Disease associations of genes located near human accelerated regions (HARs). **(A)** Heatmap showing presence (cyan) or absence (gray) of DECIPHER-reported clinical diagnoses linked to genes intersecting or proximal to HARs. Rows correspond to specific genetic disorders; columns represent individual HAR–gene pairs. **(B)** Venn diagram illustrating the overlap of five major neurodevelopmental phenotypes (Intellectual disability, Global developmental delay, Delayed speech and language development, Short stature, Microcephaly). The central region (n = 44) represents genes associated with all five phenotypic categories.

The resulting map illustrates that a subset of HAR-associated genes is linked to well-characterized hereditary disorders—primarily those affecting neurodevelopment, skeletal formation, and connective tissue. In particular, *ZMYND11*, *CNTNAP2*, *RBFOX1*, *DMD*, *HTX1*, and several X-linked loci show strong associations with syndromes involving intellectual disability, abnormalities in nervous system development, and congenital skeletal and connective tissue dysplasias.

To further evaluate the clinical relevance of HAR-associated genes, we investigated the frequency of major phenotypic categories most commonly reported among DECIPHER patients: Intellectual disability, Global developmental delay, Delayed speech and language development, Short stature, and Microcephaly.

The Venn diagram (Fig. 4B) shows the extent of overlap between these phenotypes across all genes located near HARs. The largest segment corresponds to 44 genes associated with all five categories simultaneously. Such substantial overlap suggests a high degree of shared molecular pathways underlying growth impairment, cognitive dysfunction, and nervous system maturation. Smaller clusters represent genes specific to individual phenotypes (e.g., only Microcephaly or only Delayed speech and language development), indicating potential specialization of certain HAR regulatory elements in distinct biological processes.

Overall, our findings show that many HAR-associated genes contribute to a broad spectrum of overlapping neurodevelopmental phenotypes. This supports the hypothesis that the accelerated evolution of regulatory elements in the human lineage may have targeted genes critical for brain development, cognitive function, and growth. Furthermore, we observe that the most frequently disrupted HARs in our cohort tend to interact with genes involved in various neurodevelopmental disorders, highlighting HARs as key elements for understanding the complex genotype–phenotype relationships in the context of NDDs.

### Relationship Between HARs and Genomic Instability

To conclude our study, we examined whether disruptions in human accelerated regions (HARs) contribute to genomic instability in neurodevelopmental disorders (NDDs). Recent observations in our clinical practice, together with published evidence, indicate an increasing frequency of genomic instability in affected patients and its potential role in the pathogenesis of NDDs [93–95]. This prompted us to investigate whether specific alterations involving HARs may underlie or accompany this phenomenon.

To evaluate a possible association between genomic instability and the presence of CNVs affecting HAR regions, we compared two patient groups: those with documented instability (n = 154) and those without it (n = 667). The results demonstrated borderline statistical significance (χ² = 3.90, df = 1, p = 0.048), with an odds ratio (OR) of 2.11 (95% CI: 0.99–4.48). This suggests that patients with genomic instability had approximately twice the odds of carrying HAR-disrupting CNVs. However, the confidence interval included a value close to one, indicating a high degree of uncertainty in the effect estimate.

The proportion of patients with HAR-associated CNVs remained high in both groups: 94.8% (146/154) in the instability group versus 89.7% (598/667) in the stable-genome group (Fig. 10). Despite the formally significant p-value, the small difference in percentages and the wide confidence interval around the OR indicate that the observed effect is weak and should be interpreted with caution. Further validation in larger cohorts is required to clarify whether HAR disruptions meaningfully contribute to genomic instability in NDDs.

**Figure 10.**
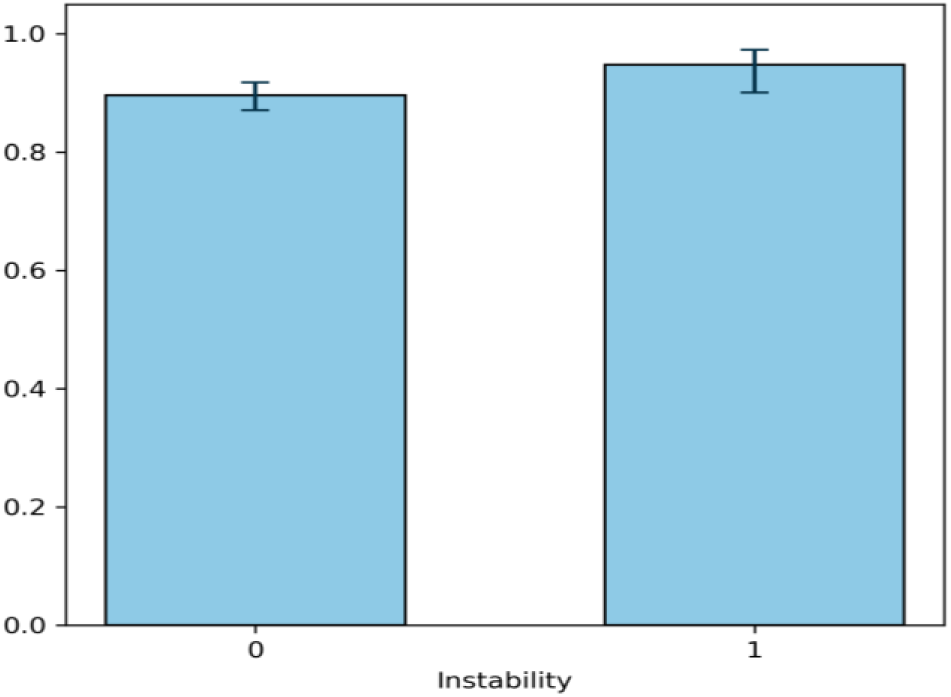
Frequency of CNV-associated HAR regions in patients with and without genomic instability. The relatively weak association between genomic instability and the presence of CNVs affecting HARs may have several explanations. First, structural variants such as CNVs tend to occur less frequently in critically important regulatory regions, as these regions exhibit a high degree of chromatin organization, elevated functional relevance, and strong deleterious consequences when disrupted. Second, cells in which such alterations do arise may undergo stronger negative selection [96], particularly when CNVs affect key enhancers that regulate brain development. Third, disruptive structural variants may lead to imbalanced expression of numerous target genes, reducing the likelihood of survival for cells harboring damage to the most “vulnerable” regulatory domains.

Thus, our results may reflect the influence of compensatory or protective mechanisms that restrict the emergence of additional structural alterations within functionally important regulatory regions, including HAR enhancers, especially under conditions of pre-existing genomic instability. Given that HAR regions play essential roles in the regulation of transcriptional programs and neural development, their disruption is likely to have disproportionately severe effects on cellular viability and organismal development. This implies that multiple interconnected protective mechanisms may act at both the molecular and cellular levels.

This interpretation is consistent with previous studies. Structural variants are significantly less likely to affect critical regulatory elements and topologically associated domain (TAD) boundaries, indicating strong purifying selection against such modifications [54, 55]. Furthermore, rare structural variants affecting genes and regulatory elements active in the brain occur at substantially lower frequencies compared with intragenic structural variants (SVs) [56]. This is largely because disruptions of essential regulatory elements in the developing brain reduce neuronal viability or result in severe neurodevelopmental phenotypes. In the context of HARs, these observations are particularly relevant: HAR regions exhibit both high evolutionary conservation and pronounced functional importance for neural development. This suggests that their disruption is likely to be strongly constrained both evolutionarily and at the level of cellular biology [15, 57].

## 3. Materials and Methods

### 3.1. Patients and Samples

Molecular karyotyping was performed to identify chromosomal abnormalities and CNV in 821 821 patients from a Russian clinical cohort presenting withм NNDs, intellectual disability, autism spectrum disorders, epilepsy, and congenital anomalies. Detailed clinical information for this cohort has been published previously [97,98]. This study was approved by the Ethics Committee of the Veltischev Research and Clinical Institute for Pediatrics of the Pirogov Russian National Research Medical University, Ministry of Health of Russian Federation, Moscow, Russia (AAAA—A18—118051590122—7 # 6, 19 June 2019). Written informed consent was obtained from the patients’ parents.

### 3.2. SNP-Array

Molecular karyotyping was performed by CytoScan HD Arrays (Affymetrix, Santa Clara, CA, USA) consisting of about 2.7 million markers. All the procedures have been repeatedly described in detail previously [99–102]. Cytogenomic variations were visualized using the Affymetrix ChAS (Chromosome Analysis Suite) software (CytoScan^®^ HD Array Version 4.1.0.90/r29400). Two human genome assemblies were used as reference sequences: GRCh37/hg19 for analyses performed before 2023, and GRCh38/hg38 for datasets generated thereafter.

### 3.3. Bioinformatic Analysis

The analysis included an annotated set of 2,737 Human Accelerated Regions (HARs) reported in Doan et al., 2016 [14]. This set was compiled by the authors based on previously published studies [6,8,9,11,103,104]. Most of these regions are located in noncoding parts of the genome (intronic or intergenic), although some intersect with promoters, noncoding RNAs, and pseudogenes. CNV coordinates obtained from the CytoScan HD platform (Affymetrix) were converted, together with HAR coordinates, to a unified genome assembly (GRCh37/hg19) using UCSC liftOver with the parameter *–minMatch=0.8* [105]. BED files were generated from CNV call tables. Overlaps between CNVs and HARs were identified using *bedtools* (v2.31.1) [106]. Annotation of intersecting HAR target genes was performed by joining CNV–HAR overlap data with the “interacting genes” sheet from Doan et al., 2016 [14] using HAR identifiers.

### 3.4. Permutation tests

To evaluate whether the observed overlaps between CNVs and HARs were statistically significant, a permutation test was performed. The observed statistic was defined as the number of CNV intervals intersecting at least one HAR. To generate a null model, CNV intervals were repeatedly randomized across the genome while preserving their length and chromosomal distribution using *bedtools shuffle* (v2.31.1) [106]. This produced random interval sets comparable in number and size to the original CNVs. For each permuted set, the number of overlaps with HARs was calculated. A total of 1,000 permutations were performed. For each analysis, a normalized z-score was computed to quantify the deviation of the observed overlap count from the permutation-derived mean. Statistical significance was assessed using an empirical p-value, defined as the proportion of permutations in which the overlap count was greater than or equal to the observed value (with a +1 correction).

### 3.5 Statistical Analysis

Statistical analysis was performed using Python 3.10 with the packages pandas (v1.5.3), numpy (v1.23.5), and scipy (v1.10.1). Absolute frequencies and percentages were used for descriptive statistics. Bivariate associations between categorical variables were assessed using Pearson’s chi-square test. Data visualization was performed using matplotlib (v3.7.1).

## Conclusions

This study represents the first large-scale analysis of the role of structural variation in human accelerated regions (HARs) in the etiology of neurodevelopmental disorders (NDDs). Based on molecular karyotyping of 821 patients, we found that CNV events affecting HAR regions are highly prevalent in this cohort, occurring in 91.7% of individuals (755 out of 821). This provides the first direct evidence that these regulatory elements—key contributors to the evolution of the human brain—are simultaneously “hotspots” of genomic instability associated with pathogenic phenotypes.

1. HARs are targets of CNVs in NDDs, with duplications experiencing stronger purifying selection. This indicates increased sensitivity of the regulatory architecture to dosage increases (copy-number gains) of enhancer elements compared with decreases. Such a pattern aligns with the concept of HARs as finely tuned regulatory modules in which excessive enhancer activity may be particularly detrimental during brain development.
2. Newly identified, previously undescribed HAR loci show high disruption frequencies. We identified three HARs with the highest CNV burden: HAR_Merge50-02702 (7.9%), HAR_Merge50-02080 (∼4.0%), and HAR_Merge50-02689 (∼2.9%). Their functional annotation links them to genes essential for neurodevelopment:

- HAR_Merge50-02702 (*PLS3*, *AGTR2*) — regulates actin cytoskeleton organization, axonal growth, synaptic plasticity, and neuroinflammatory pathways.
- HAR_Merge50-02080 (*BTNL3*, *BTNL8*) — suggests a potential connection between intestinal immune regulation (γδ T-cell signaling) and CNS development.
- HAR_Merge50-02689 (*EDA*) — affects the *EDA/EDAR/NF-κB* pathway, which regulates neuronal survival, synaptic transmission, and neuroinflammation.
3. CNVs in HARs form coordinated functional clusters rather than being randomly distributed. Cluster 1 — Adhesion cluster (3p26.3): HARs near *CHL1* and *CNTN6*, regulating axon guidance, neuronal migration, and synaptogenesis. Cluster 2 — Cross-chromosomal signaling–epigenetic module: HARs associated with regulators of key developmental pathways: *RSPO4* (Wnt), *SPRY3* (FGF), *DOC2B* (synaptic vesicle release), *ZMYND11* (epigenetic control of neuronal differentiation). This cluster represents an evolutionarily refined network coordinating growth, synaptic function, and neuronal specialization. Cluster 3 — X-linked synaptic module: HARs enriched near genes controlling synaptic architecture and plasticity: *DMD*, *IL1RAPL1*, *OPHN1*, *ZDHHC15*, *AFF2*. Its X-chromosomal localization explains the heightened vulnerability of males to NDDs and indicates that HARs fine-tune the dosage of these crucial genes.
4. HAR-associated CNVs are unique and clinically significant. Comparison with population databases (DGV, ClinGen, DECIPHER) revealed that for many HARs within these clusters, the corresponding CNVs are extremely rare or entirely absent in the general population—e.g., HARs linked to *RBFOX1*, *DOC2B*, *ZMYND11*, *AFF2*, *OPHN1*. Many of these genes also exhibit high haploinsufficiency (HI = 3), demonstrating that the CNVs observed in our cohort are not neutral polymorphisms, but pathogenic events under purifying selection.
5. HAR-associated genes display a distinct neurodevelopmental phenotype. Analysis of DECIPHER data confirmed that genes located near the most frequently disrupted HARs are significantly associated with core NDD phenotypes:

- Intellectual disability
- Global developmental delay
- Delayed speech and language development
- Microcephaly
- Short stature Strikingly, 44 genes were linked to all five phenotypes, highlighting the involvement of HAR-regulated programs in fundamental CNS growth and maturation.
6. Lack of direct correlation with genomic instability highlights the functional vulnerability of HARs. Despite the generally high burden of CNVs in HARs, their frequency did not increase in individuals with elevated genomic instability. This paradox may reflect strong compensatory mechanisms and negative selection at the cellular level: cells carrying disruptions in essential HAR enhancers may be eliminated, masking the true rate of their occurrence. This is consistent with evidence for strong depletion of structural variants in conserved regulatory elements of the brain. Our findings suggest that HARs—regulatory innovations that contributed to the expansion and functional sophistication of the human brain—are organized into coordinated functional clusters (adhesion, signaling–epigenetic, X-linked synaptic). Disruptions of these modules through CNVs lead not merely to altered expression of individual genes but to widespread imbalance across entire regulatory networks. This mechanistically explains the severity and multisystem nature of NDD phenotypes. Functionally, the most important genes include:

- *PLS3* — cytoskeleton and axonal growth
- *ZMYND11* — epigenetic neuronal programs
- *DOC2B* — synaptic neurotransmission
- X-linked regulators (*DMD*, *IL1RAPL1*, *OPHN1*, *AFF2*) — synaptic architecture This study demonstrates that CNVs in HARs represent a distinct and previously underappreciated class of enhanceropathies contributing substantially to the etiology of neurodevelopmental disorders. The identified HAR clusters and their associated target genes constitute promising candidates for functional studies, diagnostic panel development, and deeper understanding of the molecular foundations of human cognitive evolution and its vulnerabilities.

## Author Contributions

Conceptualization, M.E.I., E.D.P., I.Y.I. and E.V.S.; formal analysis, M.E.I. and E.D.P.; writing — original draft preparation, M.E.I. and E.D.P.; writing — review and editing, I.Y.I., E.V.S. and Y.A.C.; visualization, E.D.P.; resources, I.Y.I. and O.S.K.; investigation, M.E.I., E.D.P., O.S.K., E.V.G. and E.H.A.; funding acquisition, Y.A.C. and E.V.S.; supervision, I.Y.I., and Y.A.C.; project administration, E.V.S. All authors have read and agreed to the published version of the manuscript.

